# Conservation Genomic Analyses of African and Asiatic Cheetahs (*Acinonyx jubatus) Across Their Current and Historical Species Range*

**DOI:** 10.1101/2020.02.14.949081

**Authors:** Stefan Prost, Ana Paula Machado, Julia Zumbroich, Lisa Preier, Sarita Mahtani-Williams, Rene Meissner, Katerina Guschanski, Jaelle C. Brealey, Carlos Fernandes, Paul Vercammen, Luke T. B. Hunter, Alexei V. Abramov, Lena Godsall-Bottriell, Paul Bottriell, Desire Lee Dalton, Antoinette Kotze, Pamela Anna Burger

## Abstract

Cheetahs (Acinonyx *jubatus)* are majestic carnivores and the fastest land animals; yet, they are quickly heading towards an uncertain future. Threatened by habitat loss, human-interactions and illegal trafficking, there are only approximately 7,100 individuals remaining in the wild. Cheetahs used to roam large parts of Africa, and Western and Southern Asia. Today they are confined to about 9% of their original distribution. To investigate their genetic diversity and conservation status, we generated genome-wide data from historical and modern samples of all four currently recognized subspecies, along with mitochondrial DNA (mtDNA) and major histocompatibility complex (MHC) data. We found clear genetic differentiation between the sub-species, thus refuting earlier assumptions that cheetahs show only little population differentiation. Our genome-wide nuclear data indicate that cheetahs from East Africa may be more closely related to *A. j. soemmeringii* than they are to *A. j. jubatus*. This supports the need for further research on the classification of cheetah subspecies, as East African cheetahs are currently included in the Southern Africa subspecies, *A. j. jubatus*. We detected stronger inbreeding in individuals of the Critically Endangered *A. j. venaticus* (Iran) and *A. j. hecki* (Northwest Africa), and show that overall genome-wide heterozygosity in cheetah is lower than that reported for other threatened and endangered felids, such as tigers and lions. Furthermore, we show that MHC class II diversity in cheetahs is generally higher than previously reported, but still lower than in other felids. Our results provide new and important information for efficient genetic monitoring, subspecies assignments and evidence-based conservation policy decisions.

## Introduction

Cheetahs, *Acinonyx jubatus* (Schreber 1775), are currently divided into four subspecies by the Cat Classification Task Force of the IUCN Cat Specialist Group, namely *A. jubatus hecki* (Northwest Africa), *A. j. soemmeringii* (Northeast Africa), *A. j. jubatus* (Southern and East Africa) and *A. j. venaticus* (Western and Southern Asia, presently only found in Iran)^1,2^. Krausman and Morales list *A. j. raineyi* (East Africa) as a fifth subspecies^3^, but its status is under debate^1,2^. It is currently recognized as a synonym of *A. j. jubatus*, because of its close genetic relationship inferred from mitochondrial DNA^1,2^. Furthermore, there is some uncertainty on the subspecies-specific range limits of *A. j. venaticus* with speculation that it may extend into North Africa^1,4^. In 2006 the North African Region Cheetah Action Group (NARCAG) highlighted the need for genetic analyses to resolve the subspecies status of cheetahs from Algeria^4^. The most comprehensive phylogeographic analysis to date was carried out by Charruau and colleagues based on 94 cheetah samples from 18 countries (mitochondrial and microsatellite data), including regions in which cheetahs occur today or are extinct^1^. In their study, they show that *A. j. jubatus* and *A. j. raineyi* display very little genetic differentiation, a finding that was previously reported by O’Brien and colleagues^5^. They further showed that Asiatic, North-East and West African cheetahs form separate phylogenetic groups, corresponding to the currently recognized cheetah subspecies taxonomy^1^. Unfortunately, the study did not include samples from the current range of *A. j. hecki*. Schmidt-Kuentzel and colleagues reviewed the phylogeography of modern-day cheetahs and argue that the phylogeography of this subspecies cannot be assessed due to a lack of genetic data from its current range^6^. Furthermore, a recent study argues that cheetahs show very low genetic differentiation between the subspecies^7^. However, this study only included three of the four currently recognized subspecies (they did not include samples of *A. j. hecki*).

There are approximately 7,100 adult and adolescent cheetahs distributed across 33 wild sub-populations in Africa and Asia^8^ (see Fig. 1). More than half (∼60%) of wild cheetahs occur in one large population in Southern Africa^8^ (*A. j. jubatus*), while *A. j. venaticus* is represented by fewer than 50-70 only found in Iran^9^. The species’ range drastically declined over the past decades and its current extent is likely 9% of their historical distribution^8^ (2% in Asia and 13% in Africa). At the end of the nineteenth century, their distribution comprised most non-rainforest parts of Africa and much of Western and Southern Asia, from the Arabian Peninsula all the way to India, and northwards until Kazakhstan^8^. Their Asian distribution is presently limited to the central deserts of Iran^9^. The cheetah is one of the most wide-ranging carnivores with movements of up to 1,000km^10^ and home ranges up to 3,000km^2^ ^10,11^. Drivers of decline include habitat conversion and loss, persecution by pastoralists, prey declines, illegal trophy hunting, illegal trade especially as pets in the Gulf States, and armed conflicts^12–15^. The subspecies *A. j. venaticus* in Iran and *A. j. hecki* in Northwest Africa are listed as Critically Endangered by the IUCN^8,16–18^. Globally, the cheetah as a species is listed as Vulnerable^17^. However, as a majority of cheetahs occur outside formally protected areas where rates of decline are likely to be elevated, Durant and colleagues recently argued that the cheetah meets the criteria to be categorized as Endangered^8^.

**Figure 1.**
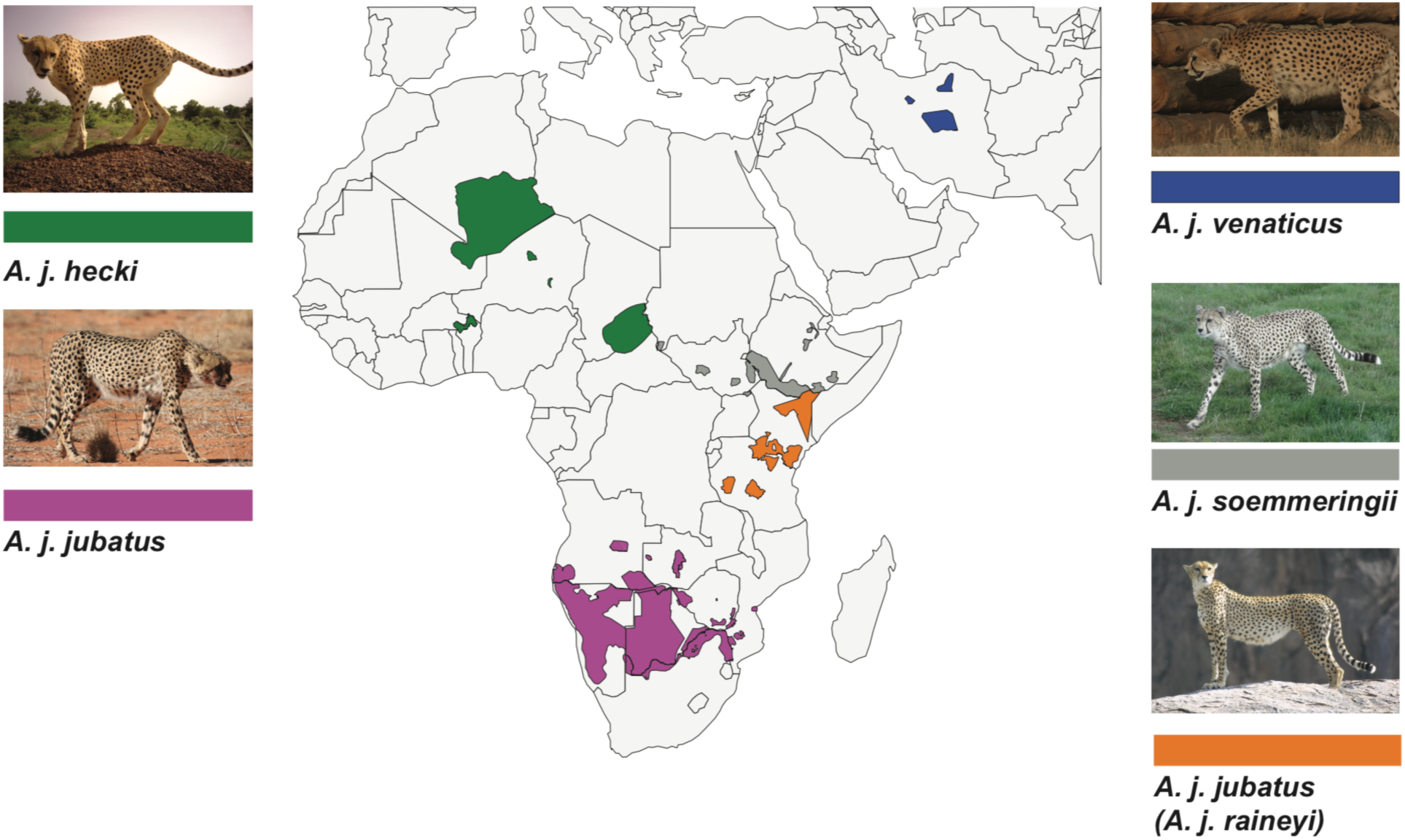
Current distribution of the five traditional cheetah subspecies (after ^3^). Green: *A. j. hecki*, purple: *A. j. jubatus*, blue: *A. j. venaticus*, grey: *A. j. soemmeringii* and orange: *A. j. raineyi*. The distribution range was adopted from the IUCN Red List^17^ and ^8^. Subspecies were assigned to the distributions using the results of ^1^ and this study. Photo credits are listed in the Acknowledgment.

In this study, we investigated genome-wide Single Nucleotide Polymorphisms (SNP), mitochondrial DNA (mtDNA) and major histocompatibility complex (MHC) class II DRB immune response gene data of the five classically recognized subspecies (*A. j. hecki, A. j. soemmeringii, A. j. jubatus* and *A. j. venaticus* and *A. j. raineyi*) to provide genomic evidence to support subspecies assignments, and to inform on-going and future conservation measures. We intend that our data can build the basis for comprehensive range-wide genetic monitoring of cheetahs through the development of reduced SNP sets. Furthermore, it can be used to guide subspecies specific conservation measures, as it sheds light on the genetic differentiation among cheetah sub-species, and will help in evidence-based decision making, e.g. for planed re-introduction projects.

## Results

To provide genomic evidence to support cheetah conservation and to better understand their phylogeographic relationships, we (i) shotgun sequenced two historical cheetah samples from Algeria and Western Sahara, (ii) generated double-digest restriction site associated DNA sequencing (ddRAD) reads for 55 modern individuals (including four parent-offspring trios, only used in the relatedness analysis), (iii) sequenced up to 929 base pairs (bp) of mtDNA for 134 modern and historical museum samples, covering wide parts of their present and historical distribution, and (iv) used amplicon sequencing to investigate MHC class II DRB exon2 haplotypes in 46 modern and historical cheetahs. For a complete sample list see Supplementary Table S1. We further downloaded genomic read data from three East African individuals from ^19^ (Genbank: SRR2737543 - SRR2737545). Here, we refer to individuals from East Africa as *A. j. raineyi* following the classical subspecies assignment for simplicity, but acknowledging that this is currently not a recognized subspecies. Our genome-wide data set thus includes individuals of each of the five classical subspecies.

### Distinct genomic differentiation and conservation status of the five classical cheetah subspecies

Subspecies or conservation unit assignments are crucial to carry out targeted conservation efforts. Therefore, we analysed genome-wide SNP data (3,743 SNPs after filtering) for 46 individuals of the five classical recognized cheetah subspecies (see Supplementary Table S1; including the three individuals from ^19^). On the contrary to the current and in line with the classical subspecies taxonomy, the principal component analyses (PCA) supported a genomic differentiation into five clusters (Fig. 2A and Supplementary Fig. S1). The five clusters correspond to the four currently recognized subspecies (*A. j. jubatus, A. j. soemmeringii, A. j. hecki and A. j. venaticus*) and *A. j. raineyi*. This is further confirmed by admixture analyses (Fig. 2B and Supplementary Fig. S2). Analyses of different subsets support a separation of the five subspecies with little to no signatures of admixture (Fig. 2B and Supplementary Fig. S2). We detected limited admixture between *A. j. raineyi* and either *A. j. soemmeringii* or *A. j. jubatus*. We further ran a separate admixture analysis including all individuals of *A. j. soemmeringii* and *A. j. jubatus*. We did not detect any signatures of admixture between the two subspecies (Supplementary Fig. S3). Genetic distances, measured using *F*ST, were the highest (0.49696) between the two endangered subspecies, *A. j. hecki* and *A. j. venaticus*, and lowest (0.15660) between *A. j. soemmeringii* and *A. j. raineyi* (Table 1). Currently, there is little known about the evolutionary history of the genus *Acinonyx* (summarized in ^20^). We thus reconstructed a maximum likelihood tree based on genetic distances from the SNP data (Fig. 2C) using the puma (*Puma concolor*) as outgroup. The first divergence event separates Asiatic and African cheetahs (with a bootstrap support of 74). We found each subspecies to form monophyletic clades with high support (bootstrap support: 90-100; Fig. 2C). The only subspecies with lower support was *A. j. hecki* (bootstrap support: 70). However, the splits separating the African subspecies show lower bootstrap support values (48-90). Individuals of *A. j. raineyi* form the sister clade to *A. j. soemmeringii* (bootstrap support: 90).

**Table 1:**
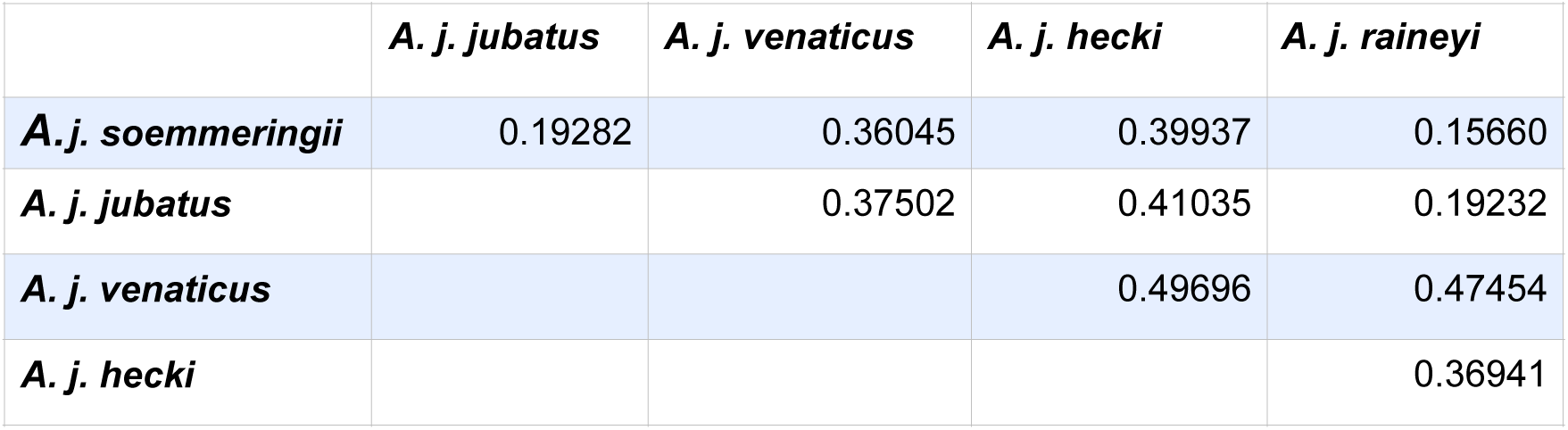
*F*_ST_ values between the five traditional subspecies of cheetah.

|  | *A. j. jubatus* | *A. j. venaticus* | *A. j. hecki* | *A. j. raineyi* |
| --- | --- | --- | --- | --- |
| *A. j. soemmeringii* | 0.19282 | 0.36045 | 0.39937 | 0.15660 |
| *A. j. jubatus* |  | 0.37502 | 0.41035 | 0.19232 |
| *A. j. venaticus* |  |  | 0.49696 | 0.47454 |
| *A. j. hecki* |  |  |  | 0.36941 |

**Figure 2:**
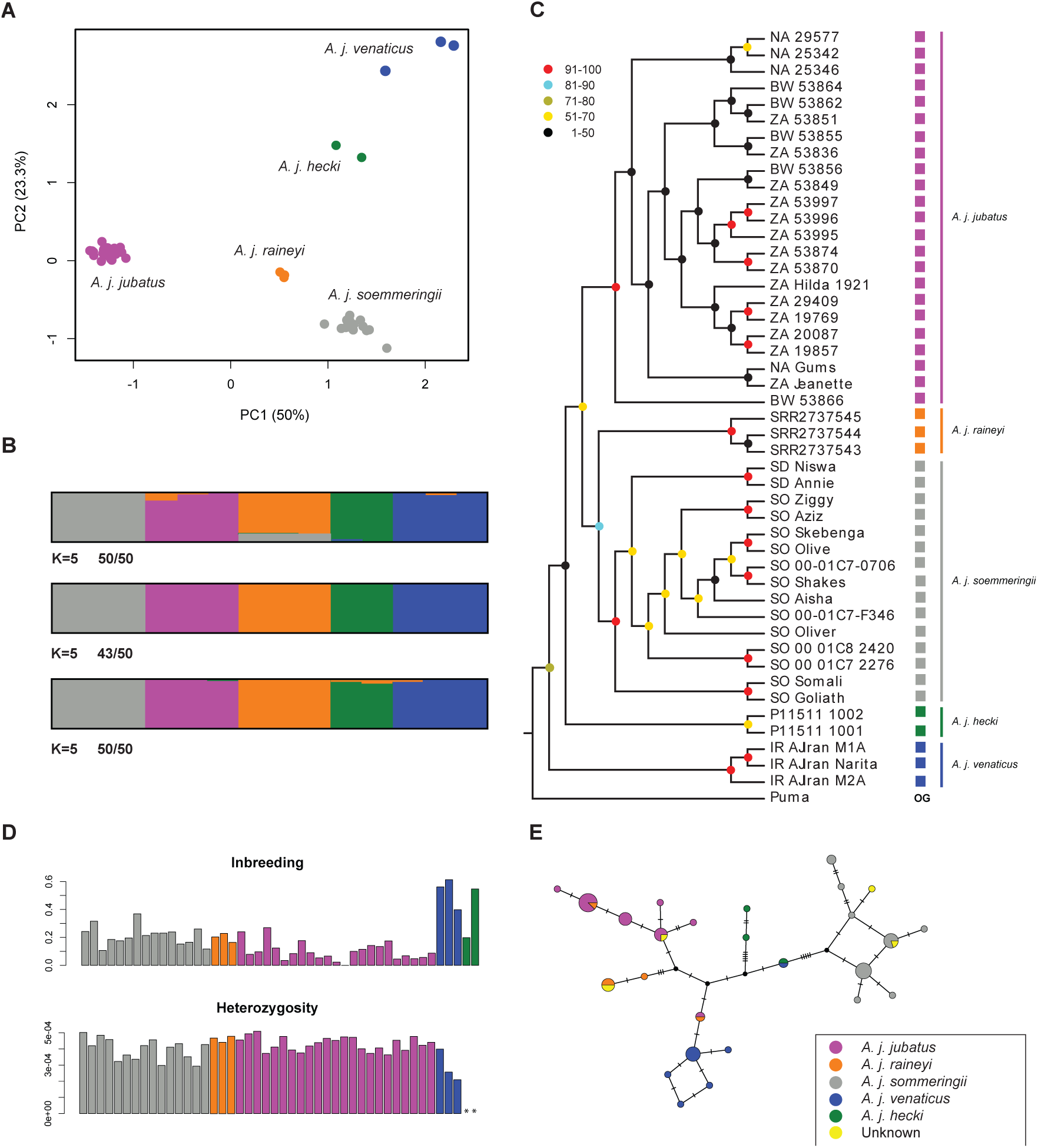
Population genomic analyses of genome-wide nuclear SNP data for 46 individuals (A-D) and mitochondrial DNA data from 58 individuals (E). (A) PCA analysis of population structure underlying the 3,743 filtered SNPs. The clustering corresponds to the morphological subspecies classification. Blue: *A. j. venaticus*, green: *A. j. hecki*, grey: *A. j. soemmeringii*, purple: *A. j. jubatus*, orange *A. j. raineyi*. (B) Admixture analyses for K=5 for the three taxon-replicates. Numbers indicate how many individual runs of the 50 replicates support this grouping. Colors as in Fig. 2A. (C) Phylogenetic relationships of representatives for the five traditional cheetah subspecies. Colors as in Fig. 2A. (D) Genome-wide inbreeding and heterozygosity indicating high inbreeding in individuals of *A. j. venaticus* and *A. j. hecki* and low heterozygosity in individuals of *A. j. venaticus*. ** indicates that individuals of *A. j. hecki* were not used in the heterozygosity analysis. Colors as in Fig. 2A. (E) Median-joining haplotype network of 929bp mtDNA. Blue: *A. j. venaticus*, green: *A. j. hecki*, grey: *A. j. soemmeringii*, purple: *A. j. jubatus*, orange *A. j. raineyi* and yellow: unknown origin.

### High inbreeding and low heterozygosity threaten the gene pool of the critically endangered Asiatic and (Northwest) African cheetahs

We carried out inbreeding analyses using two different methods, described in ^21^ and ^22^. Both inferred the highest inbreeding coefficients to be present in individuals of the two critically endangered subspecies, *A. j. venaticus* and *A. j. hecki* (Fig. 2D top panel using the method of ^22^ and Supplementary Fig. S4 using the method of ^21^), although with slightly different intensities. *A. j. jubatus* showed slightly lower inbreeding coefficients than *A. j. soemmeringii or A. j. raineyi* (Fig. 2D top panel and Supplementary Fig. S4). While most of the *A. j. jubatus* individuals are captive-bred, we do not observe any differences between these and the wild individuals. We further calculated genome-wide heterozygosity for each of the modern samples (excluding the low quality museum samples of *A. j. hecki*), which resulted in a species mean of 0.00040 (range: 0.00020 to 0.00050; Fig. 2D bottom panel). *A. j. soemmeringii* showed a mean of 0.00040 (range: 0.00029 - 0.00050), *A. j. jubatus* a mean of 0.00043 (range: 0.00036 - 0.00050), *A. j. raineyi* a mean of 0.00046 (range: 0.00044 - 0.00048) and *A. j. venaticus* a mean of 0.00029 (range: 0.00020 - 0.00040). We performed relatedness analyses to assess the impact of relatedness on our analyses. We used two different methods, due to the effects of inbreeding on the analyses^21,23^. Most *A. j. jubatus* and *A. j. soemmeringii* showed relatedness patterns (coefficient of relatedness (r = k2 / 2 + K1) and k0) indicative of 2nd to 4th generation cousins (Supplementary Fig. S5). A limited number of individuals showed sibling or parent-offspring (PO) relationships. Both methods were able to resolve relatedness for four parent-offspring trios (Supplementary Fig. S6). Comparisons between relatedness values (k2) using both methods can be found in Supplementary Fig. S7.

### Modern and historical cheetah distribution using mitochondrial DNA

The analyses of mitochondrial DNA fragments of different sizes allowed us to investigate genetic population structure throughout most of the cheetahs’ present and historical ranges, including parts of the distribution in which cheetahs are extinct today (Fig. 2E). We analysed mtDNA fragments of 681bp from 134 individuals (Supplementary Fig. S8) and 929bp from 58 individuals (Fig. 2E).

#### A. j. venaticus

We sequenced 929bp of mtDNA from different countries of *A. j. venaticus*’s past and present distribution, including Iran, Jordan, Syria, Turkmenistan, Afghanistan and India. All samples apart from one of the two Indian individuals form a single haplogroup with samples from Iran (current distribution; Fig. 2E). The other Indian individual (ID 425, see Supplementary Table S1) shares a haplotype with an individual from Chad (ID 12, *A. j. hecki*, Northwest Africa; Fig. 2E). Surprisingly, an individual from Tanzania (ID 28) and one from Zimbabwe (ID 164) show similar haplotypes to *A. j. venaticus* individuals (Fig. 2E, Supplementary Fig. S8).

#### A. j. hecki

Our sampling of *A. j. hecki* includes four individuals (from Libya, Senegal, Western Sahara and Chad) for the medium and three individuals (all but the Libyan sample) for the long mitochondrial DNA fragments. Both data sets show multiple mutations separating this subspecies from the others (Fig. 2E and Supplementary Fig. S8). All but the sample from Chad (ID 12) form a separate haplogroup.

#### A. j. soemmeringii

All individuals assigned to *A. j. soemmeringii* form a single, clearly separated haplogroup in both mitochondrial datasets (Fig. 2E, Supplementary Fig. S8). Most notably, all individuals of this subspecies show a 3bp deletion (position: 12665-12667) in the mitochondrial ND5 gene, which has previously been described in ^1^.

#### A. j. jubatus

Haplotype network reconstructions show a separate haplogroup comprising all individuals belonging to *A. j. jubatus* (apart from one samples discussed above; Fig. 2E, Supplementary Fig. S8). Four individuals from East Africa (*A. j. raineyi*) fall into the A. j. jubatus haplogroup (Fig. 2B). The same is found for the shorter mtDNA fragment.

#### A. j. raineyi

We included individuals from Tanzania, Kenia, the southern parts of Ethiopia and the southern parts of Somalia in the long mtDNA analysis. We found one Tanzanian, the Ethiopian and the Somalian sample to form a distinct haplogroup, separated from *A. j. jubatus* by 5 mutations (in 929bp; Fig. 2E). Three Tanzanian samples (from ^19^) and the Kenyan sample fell into the *A. j. jubatus* haplogroup. One Tanzanian sample showed a haplotype closely related to *A.j*. *venaticus* (discussed above). In general, we found a close relationship between haplotypes of *A. j. raineyi* and *A. j. jubatus* (Fig. 2E, Supplementary Fig. S8). Using the shorter mtDNA data set, we confirmed the presence of two different mtDNA groups in our Tanzanian and Kenyan sampling (Supplementary Fig. S8).

### Mitochondrial subspecies assignments using mini-amplicons

We tested the presence of the subspecies specific 3bp deletion in our 13 *A. j. soemmeringii* samples for which we have generated genome-wide nuclear DNA data using mini-amplicons. We found the deletion to be present in 11 individuals. The combination of all three mini-barcodes clearly put those individuals into the *A. j. soemmeringii* haplogroup. However, the other two individuals fell into the *A. j. raineyi* haplogroup (Supplementary Fig. S9).

### Adaptive immune system diversity in cheetahs

We sequenced the MHC class II DRB exon 2 of 46 individuals (belonging to four of the five traditional subspecies), which resulted in 13 nucleotide and nine amino-acid (AA) haplotypes (Fig. 3A-B, Supplementary Table S2). The most common AA haplotype was *AcjuFLA-DBR*ha16* carried by 83% of all the individuals followed by AcjuFLA-DBR*ha17 and *AcjuFLA-DBR*ha20* present in 35% and 28%, respectively (Supplementary Table S2). We found *AcjuFLA-DBR*ha21* and *AcjuFLA-DBR*ha23* present only in *A. j. jubatus*. Out of the nine AA haplotypes, we found all to be present in *A. j. jubatus* (one was only present in *A. j. raineyi)*, and five both in *A. j. soemmeringii* and *A. j. venaticus*. We found up to four different alleles to be present in single individuals (Supplementary Table S2). We inferred a haplotype diversity of 0.834 (standard deviation (std): 0.028), a nucleotide diversity (π) of 0.069 (std: 0.005) and an average of 16.2 nucleotide differences (k) for our sampling.

**Figure 3:**
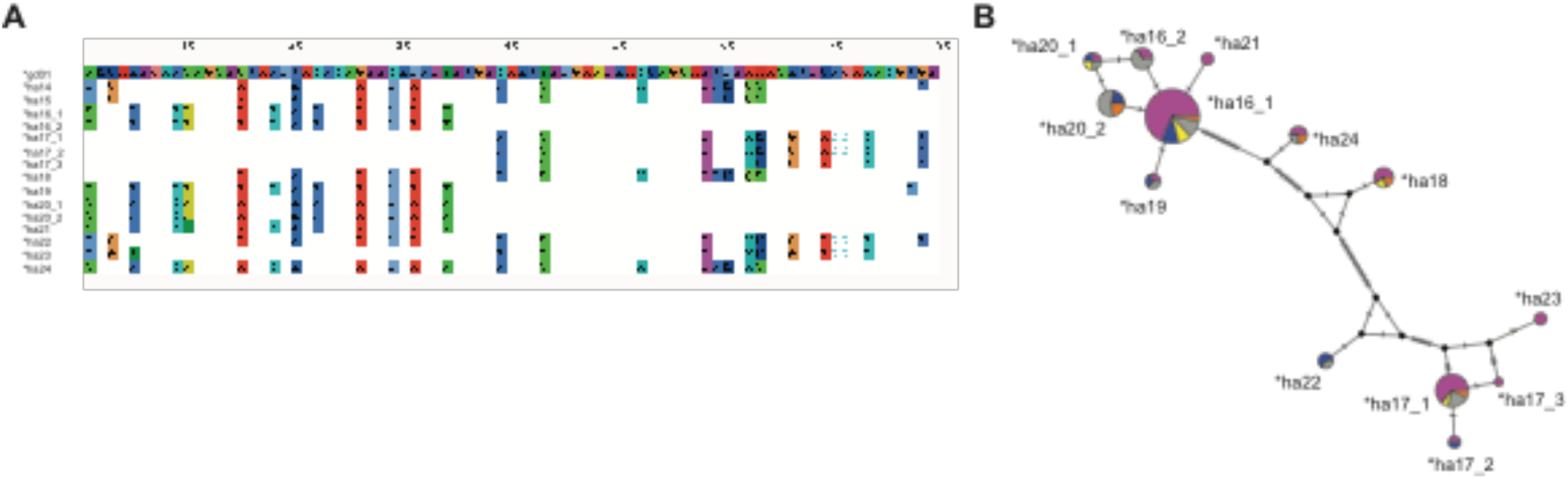
(A) Amino acid sequence and (B) a median joining network of the nucleotide sequences of the MHC II DBR exon 2. Here we abbreviated AcjuFLA-DBR*ha with *ha.

## Discussion

### Conservation status and implications

Here, we present the first genome-wide nuclear DNA support for the distinction of cheetah subspecies. PCA-based clustering, the limited and for the most part lacking admixture between subspecies, phylogenetic analyses, and high *F*ST values support five distinct groups, corresponding to the morphologically described subspecies: *A. j. jubatus, A. j. soemmeringii, A. j. venaticus, A. j. hecki* and *A. j. raineyi*. Although *F*ST differentiation between cheetah subspecies has previously been suggested to be low 7, we show that genome-wide *F*ST values (0.19 - 0.49) are comparable if not slightly higher than those of other large felids such as tigers (0.11 - 0.43 ^24^).

The cheetah as a species is classified as Vulnerable by the IUCN with a decreasing population trend^17^. Two subspecies, *A. j. venaticus* (in Iran) and *A. j. hecki* (northwest Africa) are listed as Critically Endangered by the IUCN. We found the highest levels of inbreeding in individuals of these two subspecies, further emphasizing their extremely perilous status. Furthermore, *A. j. venaticus* also showed very low genome-wide heterozygosity. We excluded our *A. j. hecki* data from this analysis, because of the low DNA quality of the museum samples. Of all the subspecies, *A. j. jubatus* showed the lowest level of inbreeding. Genome-wide heterozygosity was the highest in *A. j. jubatus* and *A. j. raineyi*. This is not surprising as *A. j. jubatus* makes up the largest continuous population of all cheetahs^8^. We have to caution that most of our *A. j. jubatus* individuals (18 out of 25) originated in captivity. However, we did not observe any differences in the genome-wide heterozygosity estimates between our captive-bred and wild caught individuals. Our genetic analyses show that cheetahs have genome-wide heterozygosity values (0.0002 to 0.0005) lower than those of other endangered big cats, such as tigers^25^ (*Panthera tigris*, 0.00049 to 0.00073; listed as Endangered) and lions^25^ (*P. leo*, 0.00048 to 0.00058; listed as Vulnerable), but higher than the Snow leopard^25^ (*P. uncia*, only one individual: 0.00023; listed as Vulnerable).

Our findings have several implications for the conservation of cheetah as well as highlighting the need for further genetic studies and monitoring.

#### (1) Subspecies specific conservation strategies

The current IUCN classification for cheetahs recognizes only four subspecies, after *A. j. raineyi* was subsumed into *A. j. jubatus*^2^. Our results based on genome-wide nuclear SNP data support the distinctions of the five classically recognized subspecies, and indicate that the merge of *A. j. raineyi* and *A. j. jubatus* might need to be revisited. Based on nuclear SNP data, *A. j. raineyi* makes up its own group with a closer relationship to *A. j. soemmeringii* than to *A. j. jubatus*. Furthermore, our analyses indicate possible admixture between *A. j. raineyi* and *A. j. soemmeringii*. Indeed, we found two individuals assigned to *A. j. soemmeringii*, using genome-wide nuclear SNP data, to carry mitochondrial haplotypes of the *A. j. raineyi* haplogroup. Interestingly, these two individuals do not show any signs of admixture (Supplementary Fig. S10). Individuals from Tanzania and Kenya showed two different mitochondrial haplogroups, one falling within the haplogroup of *A. j. jubatus* and one forming its own haplogroup, the *A. j. raineyi* haplogroup. The three individuals from Tanzania^19^, which form a separate cluster using genome-wide data, show mitochondrial haplotypes that fall within the haplogroup of *A. j. jubatus*. Genome-wide data for more individuals from East and North East Africa will be needed to fully resolve the subspecies status in this region, and to better understand the dynamics of nuclear to mitochondrial DNA in cheetahs.

Even though our genome-wide nuclear SNP data sampling from East Africa is small (n=3), our results indicate that *A. j. jubatus* and *A. j. raineyi* should be considered distinct for the purpose of management and conservation strategies. Thus, ongoing translocations of Southern African cheetahs into parts of East Africa (e.g. ^26^) may warrant closer scrutiny.

#### (2) Development of efficient range-wide genetic monitoring strategies

We generated genome-wide data for all subspecies, which can function as a baseline for the development of reduced SNP sets, e.g. for genotyping using SNP arrays or real-time PCRs enabling more cost effective and large-scale monitoring. However, more samples of the two critically endangered subspecies *A. j. venaticus* and *A. j. hecki*, and *A. j. raineyi* will be needed to avoid ascertainment bias in the selected SNP sets.

#### (3) The potential for genetic monitoring of illegal wildlife trade of cheetahs

Northeast Africa is a poaching hot-spot for the illegal cheetah pet trade, mostly to the Gulf states^15,27^. It is also likely the region with the greatest negative impact of illegal trade on wild populations of cheetahs^27^. Individuals are likely transported to the Arabian Peninsula via Somalia and Yemen. However, the origins of these animals are poorly known. Information from interdictions and interviews with traders suggest potential origins from opportunistic collections in ethnic Somali regions such as Ethiopia and Kenya^27^. Interestingly, northeast Africa is the contact zone between the two subspecies *A. j. soemmeringii* and *A. j. raineyi* (currently listed as *A. j. jubatus*). Previous studies along with our current findings indicate the presence of *A. j. soemmeringii* in South Sudan, Ethiopia and the northern parts of Somalia, and *A. j. raineyi* in Kenya, Tanzania, Uganda and the southern parts of Somalia and Ethiopia (see ^1^ and this study). Simple subspecies distinctions for illegally traded individuals and products could thus help us to quantify the respective proportion of the two subspecies in the trade, and ultimately the importance of different northeast African regions as potential countries of origin. This can form the basis for targeted programs to reduce poaching and the illegal wildlife trade of cheetahs in these countries. It will also allow evidence-based decision making for the potential release of confiscated animals into the wild including identifying appropriate potential founders for sub-specific reintroduction attempts into sites where cheetahs are currently extinct, including in Nigeria, Uganda and Rwanda. However, we have to caution that this might be complicated by possible admixture between the two, which could result in exchange of mitochondrial haplotypes. Reduced SNP set based technologies might help to overcome this potential issue.

#### (4) Environmental change and genetic diversity

Immunocompetence is an important factor for the survival of a species, and is influenced by genetic factors such as the MHC and the environment in which individuals live in ^28^. For more than two decades the cheetah has been a popular textbook example for a species with low genetic diversity, especially at MHC. Depleted immune gene diversity was previously supported by the cheetah’s ability to accept reciprocal skin grafts from unrelated individuals ^29^ and by genetic analyses (e.g. ^30^). However, these findings have been debated after ^31^ and others (e.g. ^32^) investigated allele diversity in a large sampling of cheetahs from the wild. Castro-Prieto and colleagues were able to detect a much higher genetic diversity within MHC I compared to previous studies, which they attributed to a higher sample size in their study (149 cheetahs from Namibia)^31^. However, they were not able to find any further MHC II-DRB alleles than the four previously described in ^32^. On the contrary, our sampling of 46 individuals from three subspecies (including the currently not recognized *A. j. raineyi* subspecies) resulted in nine MHC II-DRB alleles. These results are encouraging, suggesting that immunocompetence in cheetahs as coded for by the MHC may have limited impact on the conservation prospects for the species. This is consistent with the lack of substantial evidence of disease events in southern and east African wild cheetah populations which do not display evidence of compromised immunocompetence ^6,31^. Nonetheless, cheetahs show MHC II-DRB diversities lower than other large felids, such as Bengal tigers^33^ (4 alleles in 16 individuals; *P. tigris tigris*) or the Eurasian lynx in China^34^ (16 alleles in 13 individuals; *Lynx lynx*). Furthermore, Heinrich and colleagues showed that while cheetahs have lower MHC diversity than Leopards (*Panthera pardus*), they display a higher constitutive innate immunity, which could account for their relatively good health status in the wild^35^.

### Modern and historical cheetah distribution and its importance for conservation management

#### Evolutionary history

Modern cheetah appeared in Africa around 1.9 million years ago (mya)^20^. The earliest fossils of *A. jubatus* were discovered in South Africa, followed by slightly younger ones in Eastern Africa^36^. There is no information on the evolution of the cheetah subspecies based on subfossil morphological records. We thus reconstructed phylogenetic relationships using our genome-wide SNP data on modern and historical samples. We found the earliest split in the phylogenetic analysis to be that of the Asiatic and all the African subspecies (Fig. 2C). This split showed a bootstrap support of 74, the phylogenetic separation of the African subspecies showed variable bootstrap support (48-90). Each African subspecies, however, made up its own monophyletic clade with bootstrap support of 70-100. Phylogenetic relationships between the cheetah subspecies and their divergence times are heavily debated in the scientific literature (see e.g., ^1,7,37^). While our genome-wide data support the earliest split within cheetahs to be that between Asia and Africa, mtDNA analyses (e.g., ^1,37^) indicate the earliest split to be between African cheetah subspecies. This also fits phylogenetic patterns seen for example in lions ^38,39^, in which individuals from Asia are most closely related to North and Northeast African lions. However, the mitochondrial analyses of ^37^ infer a close relationship of Asiatic with Southern African (*A. j. jubatus*) cheetahs. While these results seem contradicting between the two big cat species, we have to caution that our analyses show that mitochondrial haplotypes assigned to *A. j. jubatus* can be found in East African cheetahs clearly showing a genome-wide differentiation from Southern African *A. j. jubatus*. Furthermore, estimates of the earliest divergence dates within cheetahs based on mitochondrial DNA vary strongly between analyses and range from 4.5 to 139 kya (see e.g., ^1,7,37^). Also, confidence intervals in these studies often show a high degree of overlap, making it difficult to decide on the sequence of subspecies divergences. Given the lack of a reliable mutation rate estimate for cheetah nuclear DNA, the limited bootstrap support for the underlying topology and taking the lower coverage for some of our samples into consideration, we refrained from estimating divergence times acknowledging the limitations of our data set.

#### Subspecies status of the North African cheetahs

The North African Region Cheetah Action Group^4^ (NARCAG) recommended genetic studies to identify the subspecies status for the Saharan cheetah population of Algeria, to clarify whether these individuals belong to *A. j. venaticus* or *A. j. hecki*. We included one Algerian sample in the genome-wide SNP data analyses. All analyses place this individual into the *A. j. hecki* subspecies (Fig. 2A,C,E). Mitochondrial haplotype network analyses showed a separate haplogroup for *A. j. hecki* individuals (Libya, Senegal and Western Sahara; former range states). Interestingly, the individual from Chad fell outside this haplogroup, which highlights the need for further genomic investigations of cheetahs belonging to the Chad population.

#### Distribution of A. j. venaticus

Most individuals of *A. j. venaticus* make up a clearly distinct haplogroup in the mitochondrial DNA network analyses (Fig. 2E and Supplementary Fig. S8). Interestingly, one individual from India, which was home to the *A. j. venaticus* subspecies showed a haplotype also found in Chad. It is well documented that imports of tamed hunting cheetahs from Northeaster ^40^ and Eastern Africa^41^ into India and the Arabian Peninsula were a regular occurrence during the European colonial era, which could explain this finding. Recently, Rai and colleagues proposed that cheetahs were also imported to India from Southern Africa during these times, based on a mitochondrial haplotype assigned to *A. j. jubatus*^37^. However, given the presence of *A. j. jubatus* mtDNA haplotypes in Tanzania and Kenya, we do not think this inference is currently warranted and requires more in-depth analyses. Interestingly, an individual from Tanzania and one from Zimbabwe showed similar haplotypes to *A. j. venaticus* individuals (Fig. 2E, Supplementary Fig. S8). More data, such as complete mitochondrial genomes or genome-wide SNP data will be necessary to investigate whether this is a real signal or only an artifact caused by the short length of the mtDNA fragments. Furthermore, we cannot rule out that these samples have been mislabeled sometime after their field collection. Lastly, we did not find any significant homology to nuclear mitochondrial DNA (NUMTs) copies in the published cheetah genome^19^.

#### Reintroductions of cheetahs in Asia

Several reintroduction strategies have been explored over the last years by former cheetah-range countries, including India^42^. Frequently, the reasons for animal reintroductions include conservation of the species as well as expanded tourism^43^. Ranjitsinh and Jhala identified several Indian national parks as potential candidate sites for cheetah reintroductions, though all would require extensive preparation and investment before reintroduction could be considered^42^. Genetic studies of regionally extinct individuals using museum collections are lacking, but would be crucial to assess past genetic structure in these regions and to assign individuals to their respective subspecies. Unfortunately, our sampling only included two individuals from India for the two mtDNA fragments. While one individual clustered with a sample from Chad, the other one clustered with individuals assigned to *A. j. venaticus* (which is the suspected subspecies for cheetahs from India). The two Indian individuals in ^1^ and ^37^ also showed *A. j. venaticus* haplotypes. In September 2009 Indian and International experts, at the Consultative Meeting in Gajner, suggested to introduce individuals from Africa to India^42^, as the current wild populations of *A. j. venaticus* are highly threatened and only about 50-70 individuals remain in the wild. This was further supported by a small-scale multi-locus genetic analysis^7^. In this study the authors argue that cheetah subspecies are very closely related and that genetic distances between Asian and African cheetah subspecies are equal to those within Africa, and suggested the introduction of African cheetahs to India. However, our genome-wide data shows that differentiation in cheetahs (average *F*ST of 0.34 for cheetah subspecies) is similar or even higher than that found in other large endangered felids such as the tiger^25^ (0.27), and indicate a strong genomewide differentiation of Asian and African subspecies. Based on our genome-wide data we argue against a release of African cheetahs in India, and for more genetic research to be carried out before a potential introduction of African cheetah subspecies, especially in the light of a substantial lack of information on regional adaptation in the different subspecies.

## Materials and Methods

Voucher and individual identifiers for samples used in this study can be found in Supplementary Table S1. Samples collected after 1975 were imported under the following CITES numbers: AT 16-E-0753, 16SG006329CR, 15JP001990/TE, 11US761881/9, AT 15-E-1769, D79/DFF. Additionally, we transferred samples between CITES registered institutions (see Supplementary Table S3 for the institution names and their CITES registration code).

### Laboratory Procedures Museum Samples (mtDNA)

#### DNA extraction

We followed the protocol of ^44^ for the DNA extraction from museum samples. To avoid DNA contamination we carried out all extractions in a dedicated laboratory for museum samples.

#### DNA amplification

We targeted two mitochondrial genes including 14 previously described diagnostic SNPs from ^1^ of the NADH-dehydrogenase subunit 5 (MT-ND5) and the control region (MT-CR). Four sets of primers yielding PCR products between 245 and 375 bp were used to amplify a final mtDNA fragment of 929 bp for 56 individuals and 679 bp for 134 individuals (for primer sequences see Supplementary Table S4). To avoid contamination, PCR reactions were prepared in a separate laboratory and the DNA extractions were added to the solutions in another room. Negative controls were included in each PCR reaction. The purified PCR products were sent for Sanger sequencing in both directions at Macrogen Europe Inc. (Amsterdam, Netherlands) and LCG Genomics (Berlin, Germany).

### Laboratory Procedures Mini-barcodes (mtDNA)

#### DNA amplification

We designed three mini-amplicons (see Supplementary Table S4) that amplify a total of 190bp. We then sequenced them on an Illumina iSeq 100 following the two step sequencing protocol of ^45^.

### Laboratory Procedures Museum Samples (nuclear DNA)

#### DNA extraction, Library preparation and sequencing

We extracted the DNA of two museum samples of *A. j. hecki* using the DNA extraction protocol developed by ^46^. This protocol is optimized for retaining short DNA fragments typical for highly degraded historical samples. Illumina sequencing libraries were prepared following the double-indexing strategy of ^47^. The samples were pooled in equimolar amounts and sequenced on the Illumina (San Diego, CA, USA) HiSeqX platform at SciLife, Stockholm, Sweden.

### Laboratory Procedures Modern Samples (nuclear DNA)

#### DNA extraction

We extracted DNA from 21 modern cheetah and one *Puma concolor* tissue sample using the Quiagen DNeasy Blood and Tissue kit (Qiagen, Venlo, Netherlands) and 32 diluted blood samples using the innuPREP Blood Kit (Analytik Jena AG, Jena, Germany).

#### Modern Samples - Double-digest RAD sequencing: library preparation and sequencing

Double-digest RAD sequencing was carried out for 53 samples, including the outgroup species *Puma concolor* by IGA Technologies, Udine, Italy. In brief, *in silico* analysis of the best combination of two restriction enzymes was carried out using Simrad^48^ and the Genbank aciJub1 cheetah genome assembly (GCA_001443585.1^19^). The double-digestion and library preparation was carried out following^49^, using the SphI and HindIII enzymes. Sequencing was carried out on the HiSeq2500 instrument (Illumina, San Diego, CA, USA) using V4 chemistry and paired end 125bp mode.

#### Data processing and Analyses (nuclear DNA)

First, we assessed the raw read quality was using FastQC (https://www.bioinformatics.babra-ham.ac.uk/projects/fastqc/). Next, we mapped the read data against the Aci_jub_2 cheetah genome assembly (GCA_003709585.1^19^) using BWA mem^50^ (version 0.7.17-r1188) and processed the resulting mapping files using samtools^51^ (version 1.9). We subsequently assessed the mapping quality using Qualimap^52^. We carried out adapter-trimming and duplicate removal for the two museum samples using AdapterRemoval2^53^ and Picard (https://broadinstitute.github.io/picard/), respectively. The two samples showed average coverages of 3.8x and 4.9x, respectively, after mapping and processing using BWA mem^50^ (version 0.7.17-r1188) and samtools^51^ (version 1.9).

The resulting mapping files were then processed using ANGSD^54^, which was specifically developed for population genomic analyses of low coverage data. First, we carried out filtering using SNPcleaner^22^ (v 2.24). To do so, we created a single diploid consensus sequence for all samples using samtools^51^ (version 1.9). We then filtered the resulting vcf file for (a) the presence of no more than 25% of missing data for individual sites, (b) a maximum coverage of 5,000 to avoid calling sites in highly repetitive regions, and (c) a minimum coverage of 3x for each individual. We then extracted the first two columns of the resulting bed file to generate the filtered SNP files for the subsequent ANGSD analyses.

We included 46 individuals in the population genomic analyses. First, we carried out principal component analyses using ANGSD and pcangsd^55^. Next, we looked for signatures of admixture using ANGSD and ngsAdmix^56^. In order to avoid sampling biases, due to the different numbers of samples per subspecies, we generated different sets of three randomly chosen individuals for each subspecies (except for *A. j. venaticus* and *A. j. hecki* where we only had three and two in our sampling, respectively). We further carried out a separate ngsAdmix analysis restricted to all individuals of the two subspecies *A. j. soemmeringii* and *A. j. jubatus*. We carried out 50 replicates for all ngsAdmix run ranging from k=2 to k=5. The results were analysed and visualized using CLUPMACK57. In order to investigate subspecies differentiation, we carried out *F*ST analyses using ANGSD (realSFS). Phylogenetic analyses were carried out using a combination of ANGSD, ngsDist^22^ and FastME^58^. Inbreeding was investigated using ngsF^22^ and ngsRelate^21^ (v2). We further used ngsRelate (v1^23^ and v2^21^) to carry out relatedness analyses. In order to check its performance, we first ran the tool using data from four parent-offspring trios. Lastly, we carried out heterozygosity analyses using ANGSD and realSFS (part of the ANGSD package). Here, we did not restrict our analyses to filtered sites, to be able to compare our estimates to published genome-wide heterozygosity values of other felids. Due to low coverage and quality, typical for degraded museum DNA, we removed the two museum samples from this analysis.

### Mitochondrial DNA data

For the mitochondrial DNA data we aligned all sequences using Codon Code Aligner v3.0.2 (Codon Code Corporation). We obtained the reference sequence for the cheetah mitochondrial genome from GenBank (accession number NC_005212.1^59^). We carried out parallel analyses on the two concatenated datasets that differed in the length of the fragment and the number of individuals (Supplementary Table S1). The first analysis comprised the largest mtDNA fragment of 929 bp amplified in 58 individuals. The second line of analysis incorporated the alignment of 78 individuals from ^1^ and 57 generated in this study. Median-joining networks were created using the freely available software tool, Popart^60^. We also generated median joining networks for the mini-barcode data. Furthermore, we investigated the homology of the mtDNA fragments to known NUMTs in the published cheetah genome (GCA_003709585.1^19^) using Blast.

### MHC DNA data

#### DNA extraction, Amplification and Sequencing

We used the Qiagen DNeasy Blood and Tissue kit for DNA extraction from hair and tissue samples, and the VWR PeqGold™ Tissue DNA Mini Kit Plus for blood samples. We carried out Polymerase Chain Reaction (PCR) as described in ^31^ using the primers: DRB_SL-F (GCGTCAGTGTCTTCCAGGAG) and DRB_SL-R (GGGACCCAGTCTCTGTCTCA). Indexing, multiplexing and sequencing was carried out following the Illumina Nextera XT DNA LibraryPrep reference guide. Sequencing was carried out on an Illumina MiSeq (2×250bp). We then mapped the Illumina sequencing reads against a reference Sanger sequence using BWA mem^50^ (version 0.7.11) and further processed the mapping file using samtools^51^ (version 1.3.1). We called variants using Picard v. 2.8.2 (http://broadinstitute.github.io/picard) and GATK^61^ (v.3.1.8). The main alleles were phased using the FastaAlternateReferenceMaker command in GATK, based on a minimum of 6 reads per sample, and verified manually by visualization of the coordinate-sorted BAM files using IGViewer^62^. We estimated haplotype diversity (Hd) and nucleotide diversity (π) using DNAsp^63^.

## Acknowledgements

For assistance with samples, we are very grateful to the Iranian Department of Environment, the CACP (Conservation of Asiatic Cheetah Project) and especially its former directors, Houman Jowkar, Alireza Jourabchian, Akbar Hamedanian and Hooshang Ziae. Thanks also to Stephane Ostrowski and Chris Walzer (WCS) for their assistance in Iran. We thank the curators of the Natural History Museums of London, Vienna, Berlin, Paris, Copenhagen, Stuttgart and Senckenberg, the Alexander Koenig Research Museum, the Harrison Institute, the Royal Museum of Scotland and the Royal Museum for Central Africa (Tervuren), and the Zoological Institute of Russian Academy of Sciences (AAAA–A19– 119032590102-7). We appreciate the support from the veterinarians and keepers of DECAN Rescue Centre Djibouti, Vienna Zoo, Salzburg Zoo, La Palmyre Zoo and the Zoological Garden of Montpellier. The authors further acknowledge support from the National Genomics Infrastructure in Stockholm funded by Science for Life Laboratory, the Knut and Alice Wallenberg Foundation and the Swedish Research Council, and SNIC/ Uppsala Multidisciplinary Center for Advanced Computational Science for assistance with massively parallel sequencing and access to the UPPMAX computational infrastructure. The authors also thank Henrique Leitão and Thijs Hofstede for help processing the museum specimens in the aDNA lab. We thank Christine and Urs Breitenmoser, Sven Winter and Menno de-Jong for their helpful comments on the manuscript. PB acknowledges funding from the Austrian Science Foundation FWF P29623-B25. Photo credits: *A. j. hecki* (©ZSL-RWCP/Panthera/IUCN Cat SG in collaboration with APN), *A. j. jubatus* (© Sarah Durant), *A. j. venaticus* (©Sarah Durant), *A. j. soemmeringii* (© Mark Holden) and *A. j. raineyi* (© Sarah Durant).

## SUPPLEMENTARY FIGURES AND TABLES

**Supplementary Figure S1: PCA analyses for the genome-wide SNP data (PC1 versus PC3)**. Blue: *A. j. venaticus*, green: *A. j. hecki*, grey: *A. j. soemmeringii*, purple: *A. j. jubatus*, orange: *A. j. raineyi*.

**Supplementary Figure S2: Admixture results for the three independent replicate runs**. Results are shown for 50 replicates of K=2, K=3, K=4 and K=5. Each run used different individuals for *A. j. jubatus* and *A. j. soemmeringii*. Only groupings supported by more than four replicates are shown.

**Supplementary Figure S3: Admixture results for the *A. j. jubatus* and *A. j. soemmeringii* dataset**. Results are shown for 50 replicates of K=2, K=3, K=4 and K=5. Only groupings supported by more than four replicates are shown.

**Supplementary Figure S4: Inbreeding values calculated using the method of** ^**1**^. Blue: *A. j. venaticus*, green: *A. j. hecki*, grey: *A. j. soemmeringii*, purple: *A. j. jubatus*, orange: *A. j. raineyi*.

**Supplementary Figure S5: Plot showing K0 versus the coefficient of relatedness (r) for *A. j. jubatus* and *A. j. soemmeringii***. The expected relatedness are shown for different areas in the plot. PO…parent offspring, Sib…sibling, 2nd…second generation cousins, 3rd…third generation cousins, 4th…fourth generation cousins and UR…unrelated. Grey: *A. j. soemmeringii* and purple: *A. j. jubatus*.

**Supplementary Figure S6: Relatedness (K2) of four parent-offspring trios**. Top panel: relatedness inferred without accounting for inbreeding, and bottom panel relatedness inferred after correcting for inbreeding. Dark blue indicates a close relationship and a white no relationship between individual pairs.

**Supplementary Figure S7: Relatedness (K2) found in the 43 cheetah samples**. Top panel: relatedness inferred without accounting for inbreeding, and bottom panel relatedness inferred after correcting for inbreeding. Dark blue indicates a close relationship and a white no relationship between individual pairs.

**Supplementary Figure S8: Median-joining haploptype network of 679 bp mitochondrial DNA**. Blue: *A. j. venaticus*, green: *A. j. hecki*, grey: *A. j. soemmeringii*, purple: *A. j. jubatus*, orange: *A. j. raineyi*.

**Supplementary Figure S9: Median-joining haploptype network of 190 bp mitochondrial DNA**. Blue: *A. j. venaticus*, green: *A. j. hecki*, grey: *A. j. soemmeringii*, light grey: *A. j. soemmeringii* from the ddRAD analysis, purple: *A. j. jubatus*, orange: *A. j. raineyi*. We removed samples with questionable or unknown origin for this analysis.

**Supplementary Figure S10: Admixture results for the run including the two *A. j. soemmeringii* ddRAD samples (*) that showed mitochondrial haplotypes in the *A. j. raineyi* haplogroup**.

**Table S1: Samples used in this study**.

**Table S2: MHC II DBR exon 2 haplotypes and their frequency in the different subspecies**.

**Table S3: CITES registered institutions and their registration numbers**. Samples were transported between institutions under the listed CITES registration numbers.

**Table S4: Characteristics of the primers employed in this study**. aMT indicates the primers used for highly fragmented museum samples.

