## Supplementary figures and images for "Conservation Genomic Analyses of African and Asiatic Cheetahs (*Acinonyx jubatus) Across Their Current and Historical Species Range*"

### Supplementary Figure 1

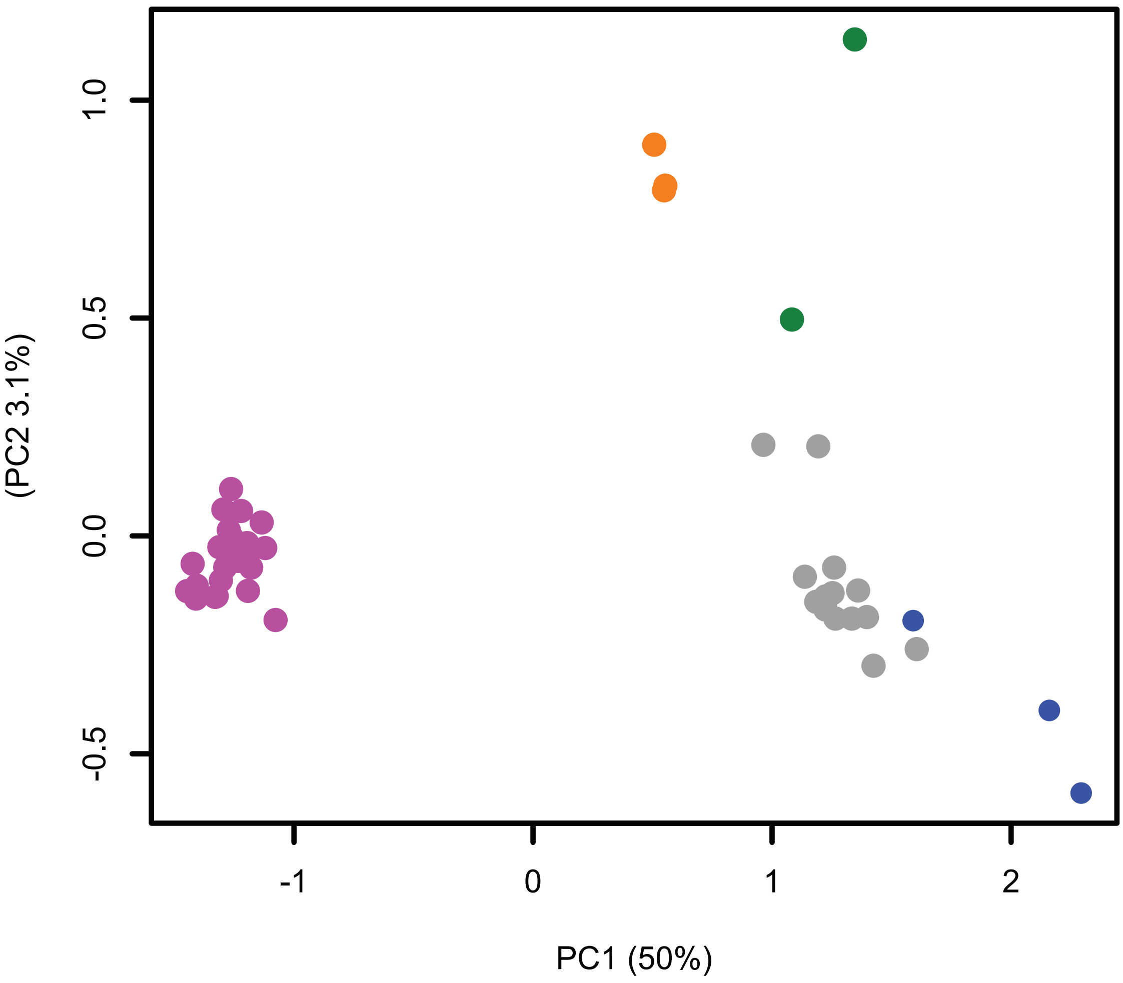

### Supplementary Figure 2

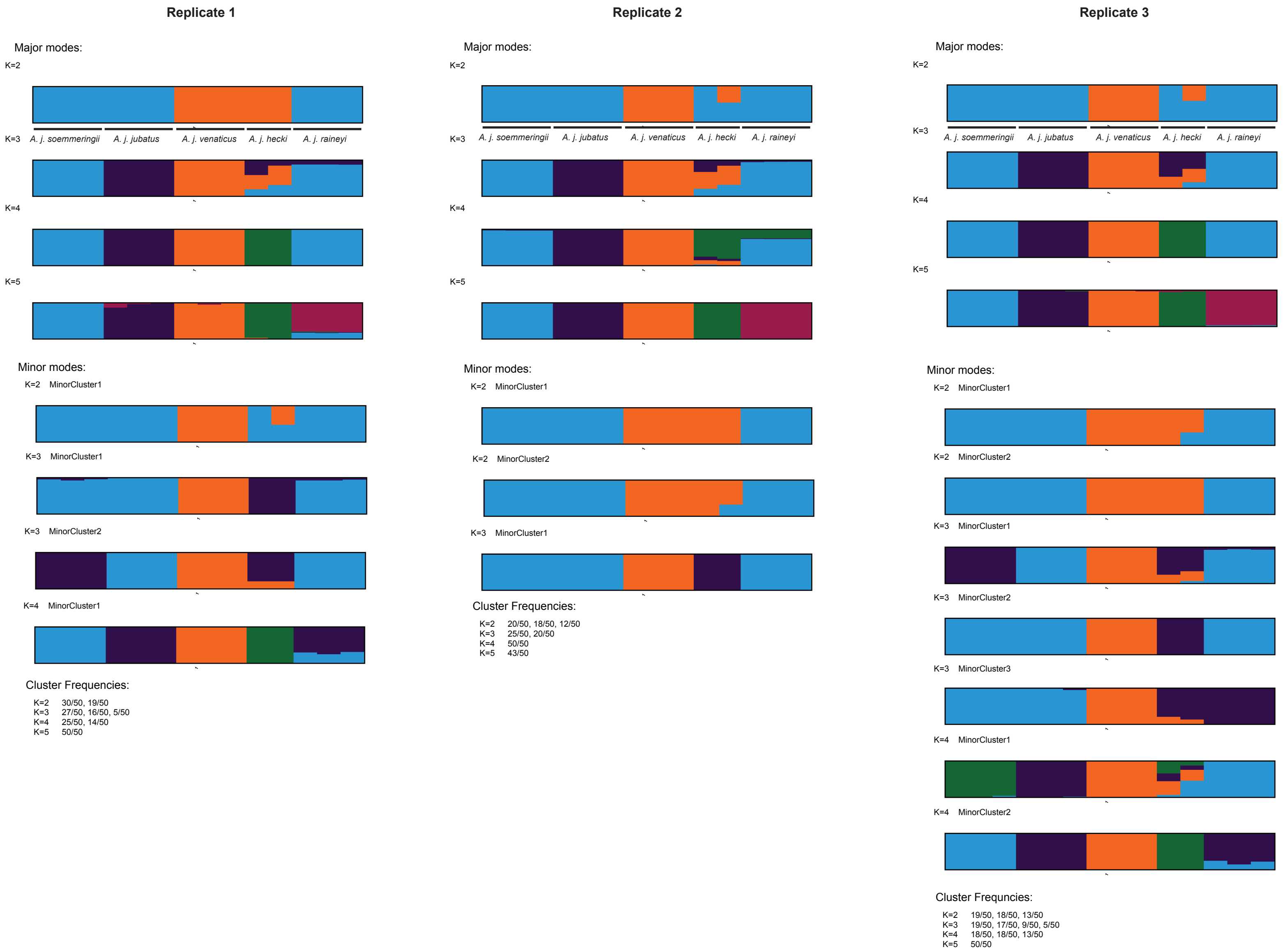

### Supplementary Figure 4

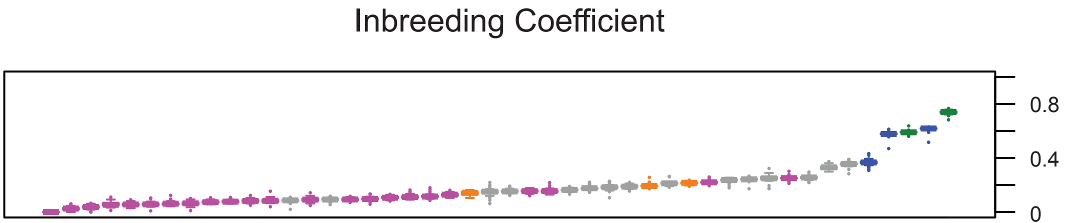

### Supplementary Figure 5

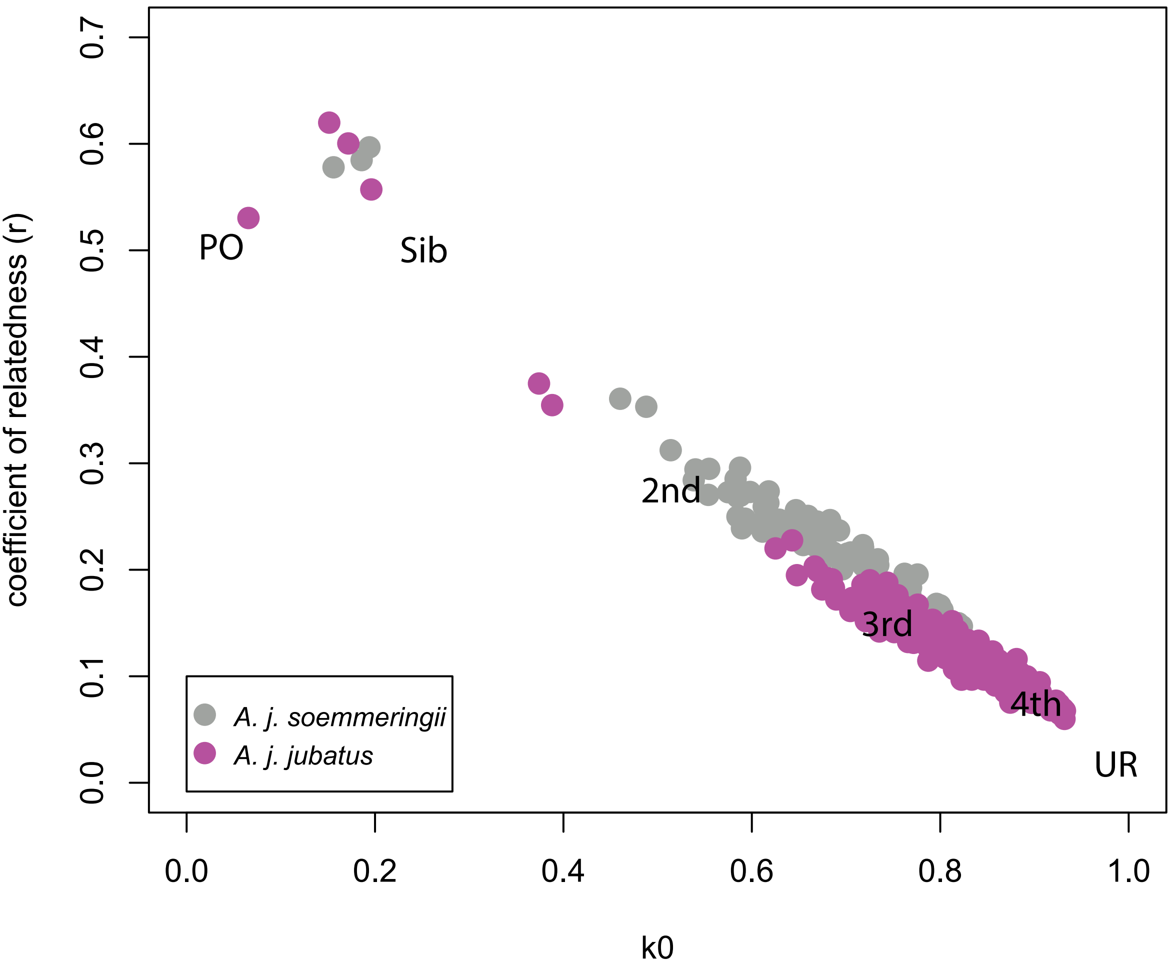

### Supplementary Figure 6

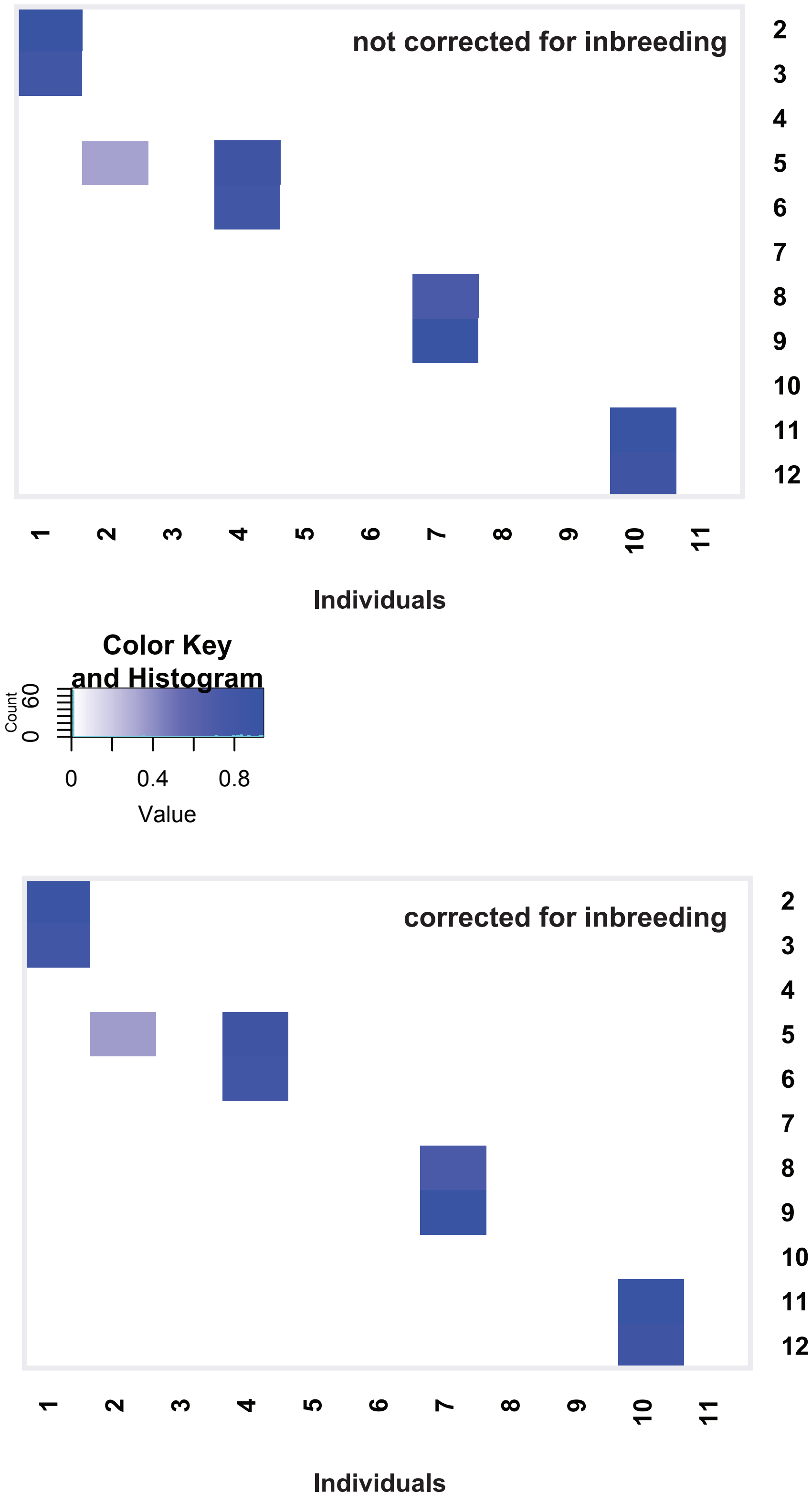

### Supplementary Figure 7

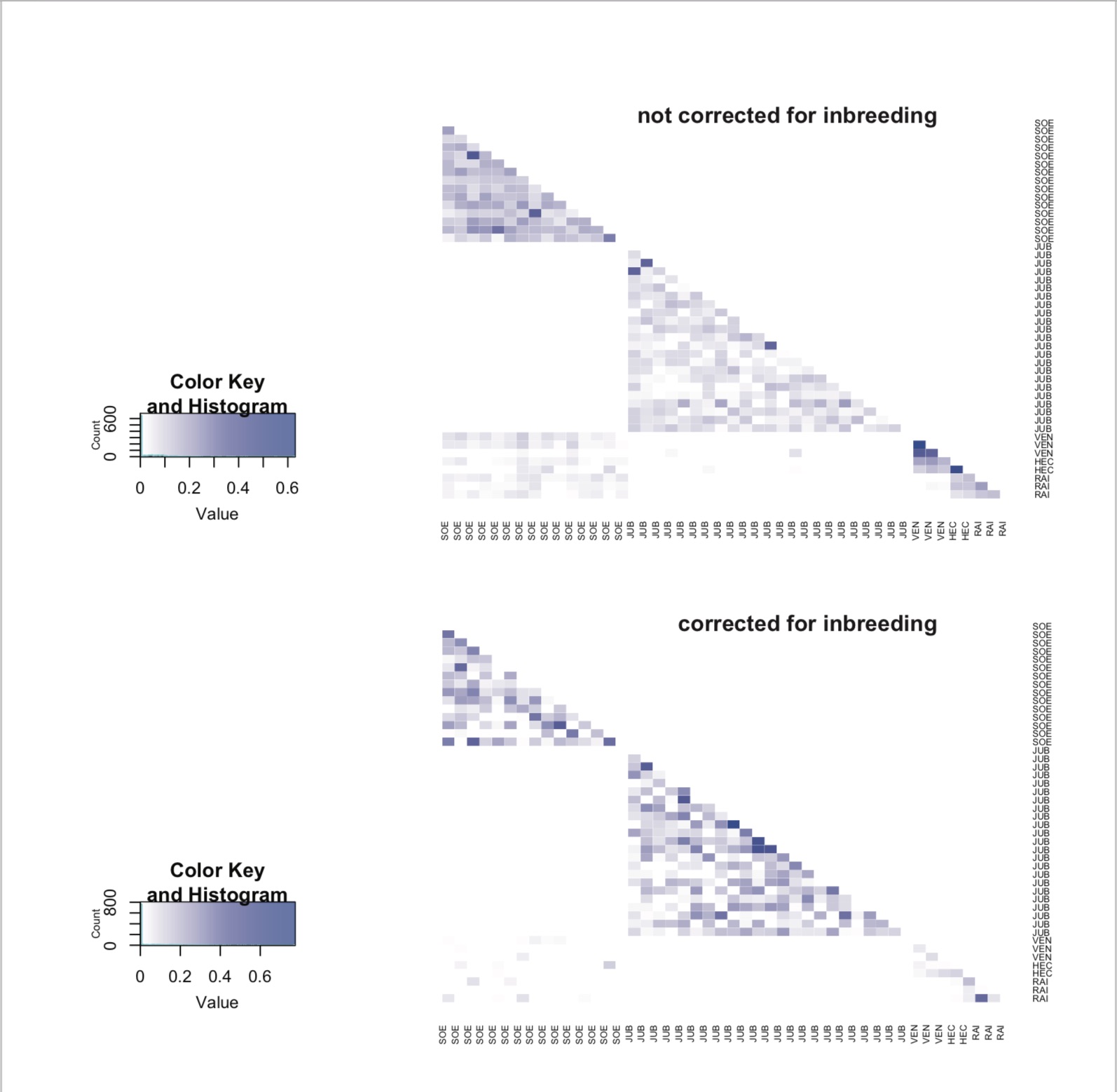

### Supplementary Figure 8

#### Medium Size (679bp)

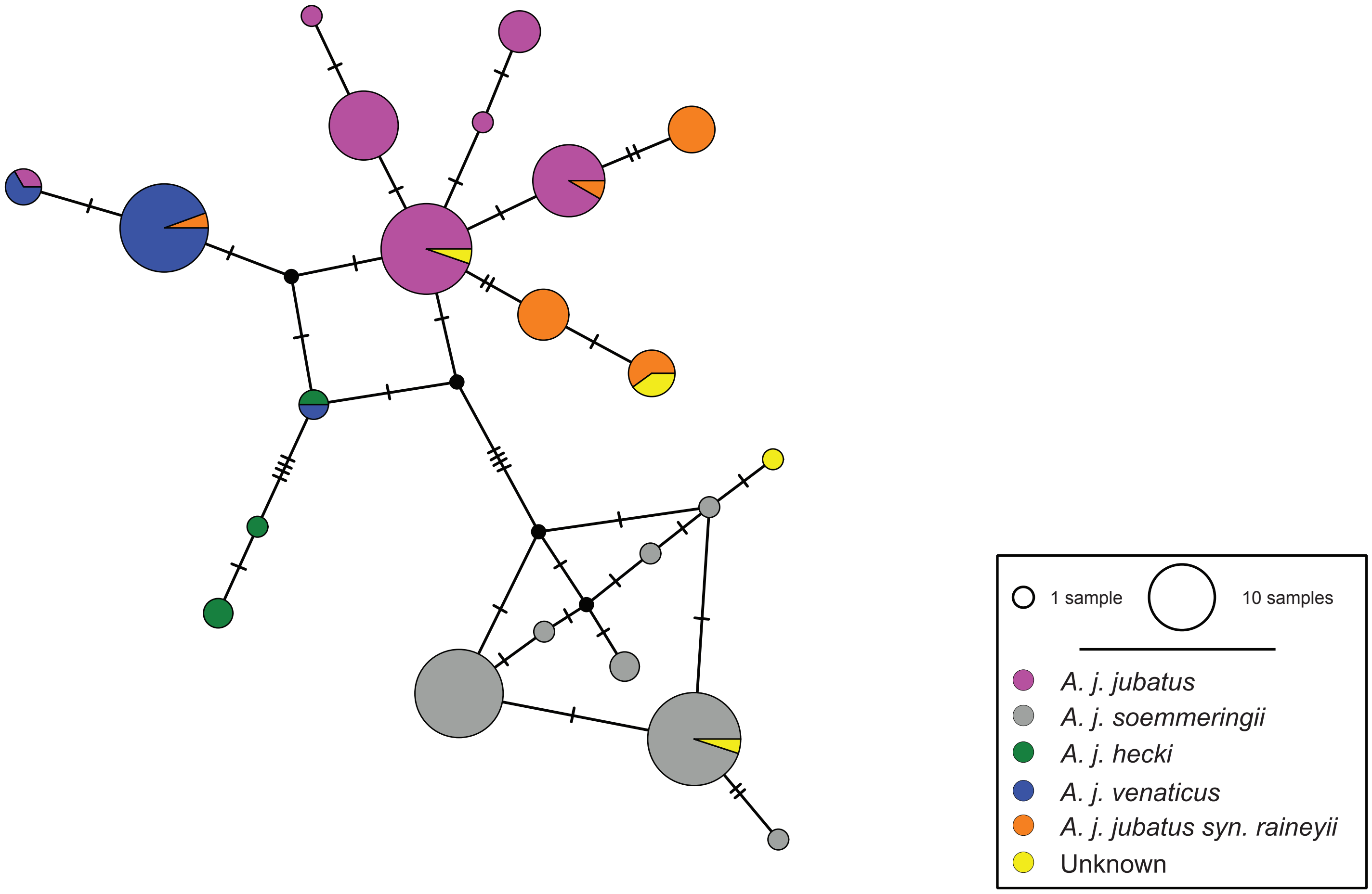

### Supplementary Figure 9

#### Mini-amplicons (190bp)

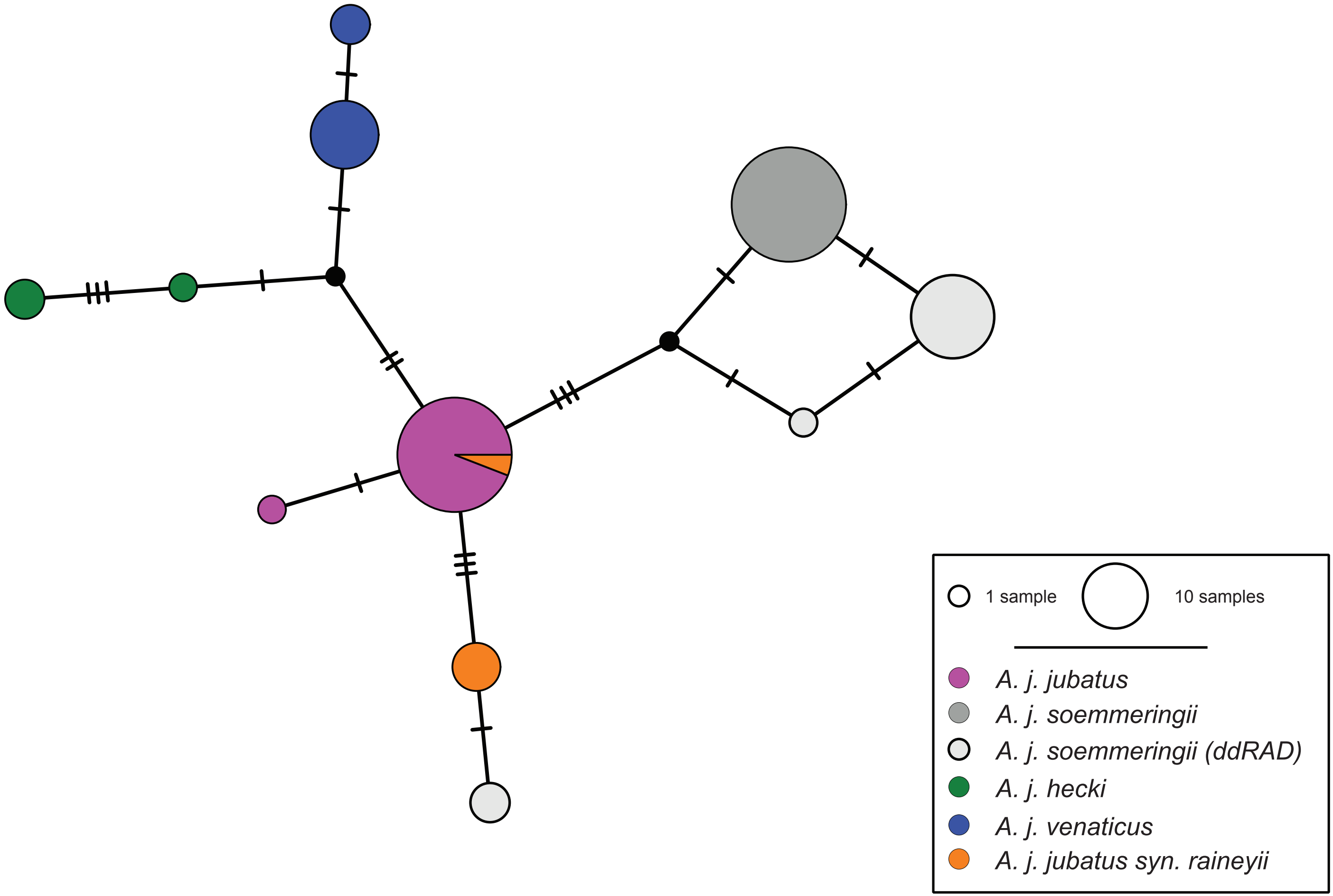

### Supplementary Figure 10

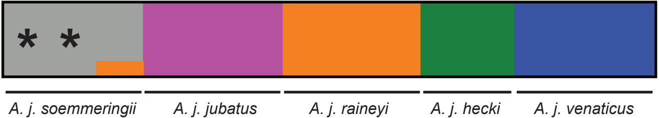
