## Supplementary Figure 3 for "Conservation Genomic Analyses of African and Asiatic Cheetahs (*Acinonyx jubatus) Across Their Current and Historical Species Range*"

### Major modes:

K=2

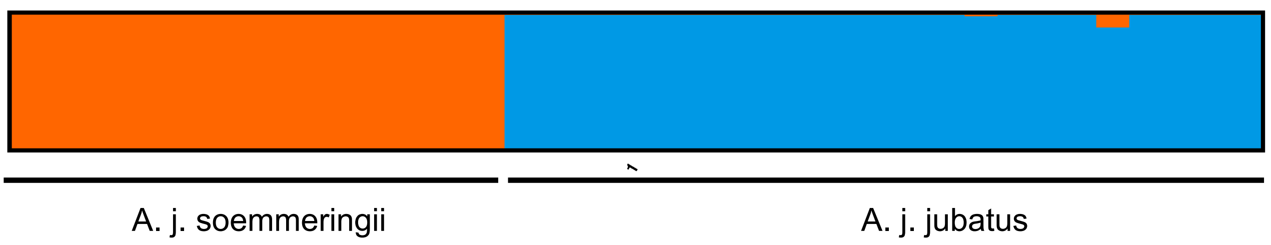


K=3

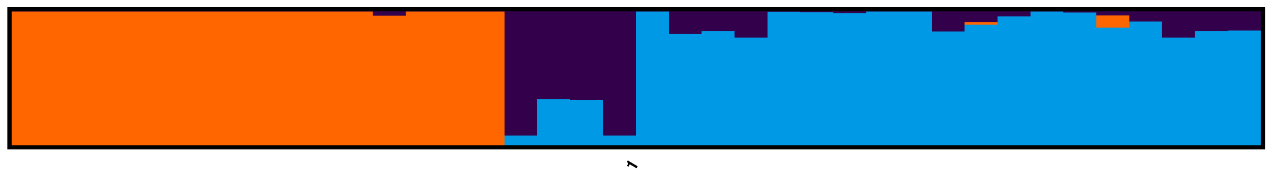

K=4

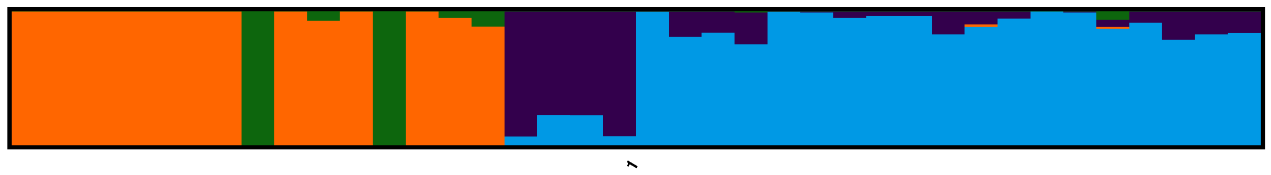

K=5

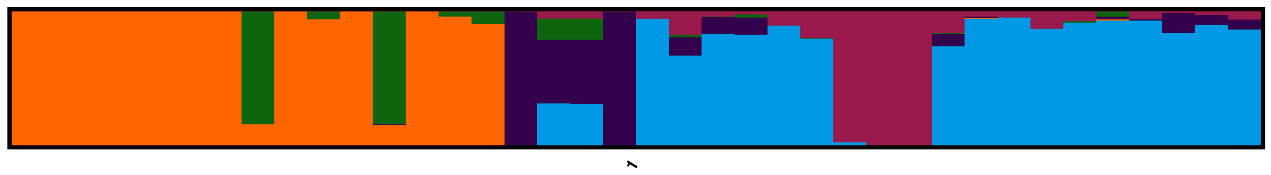

### Minor modes:

K=3 MinorCluster1

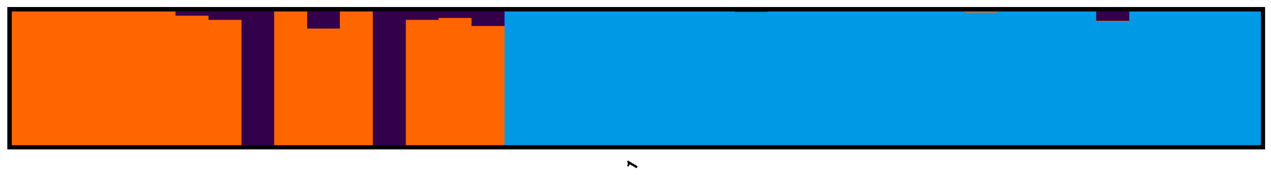

K=4 MinorCluster1

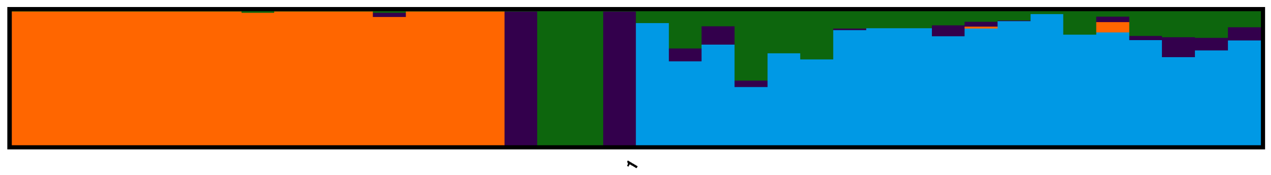

K=5 MinorCluster1

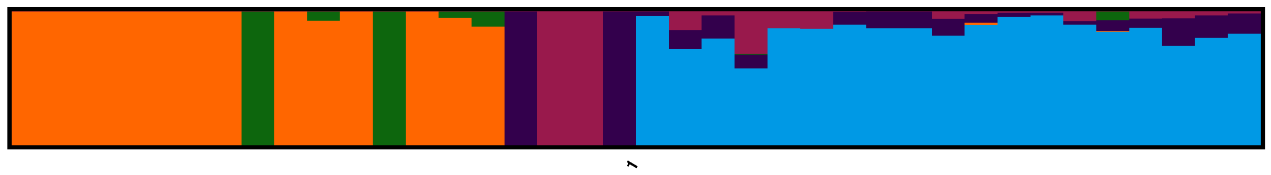

K=5 MinorCluster2

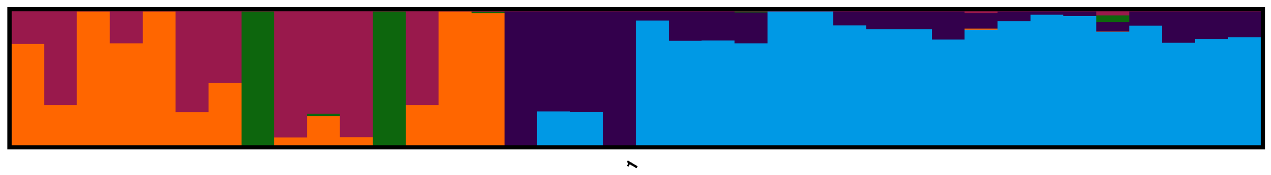

### Cluster Frequencies:

K=2 50/50
K=3 27/50, 16/50
K=4 29/50, 8/50
K=5 19/50, 16/50, 15/50
